# Hidden Markov models detect recombination and ancestry of SARS-CoV-2

**DOI:** 10.1101/2025.11.08.687354

**Authors:** Nobuaki Masaki, Trevor Bedford

**Author notes:** Corresponding author: Nobuaki Masaki, MRC Laboratory of Medical Sciences, London, UK.

## Abstract

When individuals are co-infected with distinct SARS-CoV-2 lineages, homologous recombination can generate mosaic genomes carrying mutations from both parental lineages. A variety of methods exist to detect recombinant sequences and their parental lineages in surveillance-scale datasets comprised of millions of SARS-CoV-2 genomes. However, these methods often rely on user-specified parameters, such as the probability a recombination breakpoint occurs between adjacent positions on the query sequence. In this study, we devise a hidden Markov model that detects recombinant SARS-CoV-2 sequences and identifies their parental lineages within a test set of sequences. Our method does not depend on user-specified parameters and can accommodate de novo mutations on the query sequence that are not present in the predicted parental lineages. To achieve this, we use maximum likelihood to estimate parameters that characterize the transition and emission probabilities in our hidden Markov model. Applying our method to 440,307 SARS-CoV-2 sequences sampled in England between September 2020 and March 2024, we detect 7,619 recombinant sequences corresponding to 1.73% (95% CI: [1.69%, 1.77%]) of all sampled sequences. We observe a positive association between the proportion of query sequences detected as recombinant in each week and community SARS-CoV-2 prevalence. This is consistent with higher prevalence increasing the risk of co-infection by distinct lineages and promoting the emergence of recombinant sequences. Finally, we observe localized clusters of recombination breakpoints within spike and in intergenic regions.

## 1 Introduction

Recombination is thought to occur in coronaviruses via a copy-choice mechanism in which the viral RNA-dependent RNA polymerase switches template strands during negative strand synthesis (Chrisman et al. 2021). When hosts are co-infected by multiple SARS-CoV-2 lineages, this template switching results in recombinant genomes sharing genetic material from both lineages (Trémeaux et al. 2023).

One of the most notable recombinant lineages that emerged during the pandemic is XBB. Phylogenetic analysis indicates that this lineage was derived from a recombination event between two Omicron lineages (BJ.1 and BM.1.1.1) and resulted in significant reduction in neutralization from human serum samples (Tamura et al. 2023). The effective reproduction number of XBB was estimated to be 1.23 and 1.20 times higher than its parental lineages BJ.1 and BM.1.1.1, respectively, using epidemic data from late 2022 (Tamura et al. 2023). The derived lineage XBB.1.5 spread widely and reached a peak frequency of 55% globally in epidemiological week 12 of 2023 (Erkihun et al. 2024). Because recombination can combine mutations from different SARS-CoV-2 lineages that jointly confer a growth advantage to the recombinant genome, systematic surveillance and robust statistical detection of recombinant lineages are crucial. Recombination-aware analyses applied over long evolutionary timescales have also been used to investigate the evolutionary origins of SARS-CoV-2, including genomic regions such as the receptor binding domain (Lytras et al. 2022; Esquivel Gomez et al. 2024).

A wide range of computational approaches have been developed to detect recombination in viruses. Broadly, similarity methods such as SimPlot visualize how a query sequence’s similarity shifts across the genome relative to putative parental lineages (Salminen et al. 1995; Samson et al. 2022). RDP4 examines all triplets within a set of sequences and applies a suite of tests (e.g., GENECONV, MaxChi, Bootscan, 3SEQ) to detect recombination breakpoints and assign parental sequences (Sawyer 1989; Posada and Crandall 2001; Martin et al. 2015; Lam et al. 2018). However, the number of comparisons is cubic with respect to the sample size, which is infeasible for large-scale datasets.

Phylogeny-based methods such as GARD detect breakpoints by fitting independent phylogenies to alignment segments and comparing model fit across candidate partitions (Kosakovsky Pond et al. 2006). The repeated tree-fitting and model-comparison steps are computationally intensive, so GARD is generally applied to downsampled alignments instead of surveillance-scale datasets comprising millions of genomes.

More recently, SARS-CoV-2-specific tools have been designed to operate on surveillance-scale datasets. Bolotie uses a hidden Markov model (HMM) where the latent states represent SARS-CoV-2 lineages (Varabyou et al. 2021). The Viterbi algorithm is used to assign a parental lineage to each position. RIPPLES identifies candidate recombinant sequences by scanning a global mutation-annotated phylogeny for unusually long branches that may represent recombination events (Turakhia et al. 2022). For each candidate recombinant sequence, RIPPLES partitions the genome into multiple segments and re-places each onto the global phylogeny using maximum parsimony. RecombinHunt compares segment-wise mutation patterns on a query sequence to lineage-specific profiles (Alfonsi et al. 2024). It constructs a cumulative likelihood profile across the genome and uses the Akaike information criterion to choose between three models with zero, one, or two breakpoints.

Although these SARS-CoV-2–specific tools can be applied to surveillance-scale datasets, each has method-specific limitations. Bolotie’s HMM does not model de novo mutations or genotyping errors, which can result in spurious state switches when the query sequence harbors mutations absent from the mutation profile of its true lineage. The HMM’s transition probability is also user-specified, making breakpoint detection sensitive to this choice. RIPPLES relies on a mutation-annotated phylogeny. Uneven sampling and sequencing artifacts can inflate or deflate the long-branch signal used to identify candidate recombinants. Moreover, the threshold for the long-branch signal is defined by the user, and the initial candidate set of recombinant sequences is sensitive to this chosen cut-off. RecombinHunt relies on several hard evidence gates (e.g., declaring a genome non-recombinant when it differs from the most likely lineage by two or fewer mutations). Classification with these thresholds are likewise sensitive to de novo mutations and genotyping errors. Finally, both RIPPLES and RecombinHunt permit at most two breakpoints, even though recombinant lineages with more breakpoints have been detected.

In this paper, we develop a method to detect recombinant SARS-CoV-2 sequences within a test set of sequences collected over a short interval (a few days to a week). Our method employs an HMM inspired by the Li and Stephens (2003) model that accounts for de novo mutations and genotyping errors in both recombinant and non-recombinant sequences. For each test sequence, we estimate a pseudo-frequency for observing alleles absent from the true parental lineage, and the lineage-transition probability between consecutive sites. We implement an efficient version of the forward algorithm to speed up estimation of these frequencies (see Section S1 of the Supplementary Materials). We then predict the local Pango lineage ancestry, defined as the sequence of Pango lineages ancestral at each genomic position of the test sequence, using lineage-specific nucleotide frequencies computed from prior sequences. We classify a test sequence as a recombinant if the predicted local Pango lineage ancestry contains one or more lineage transitions. Our method does not rely on a phylogeny or any user-defined parameters, and can accommodate any number of breakpoints.

We evaluate performance in a simulation where we generated synthetic recombinant and control genomes from SARS-CoV-2 sequences sampled between January and March 2022. We also apply our method to 440,307 SARS-CoV-2 genomes from GenBank (Benson et al. 2013) sampled in England between September 2020 and March 2024 to identify recombinant sequences and measure their frequency through time and occurrence across parental lineage pairs.

## 2 Materials and methods

Figure 1 summarizes our workflow for detecting the local Pango lineage ancestry of SARS-CoV-2 sequences from GenBank. In this section, we describe each component of our method in detail.

**Figure 1.**
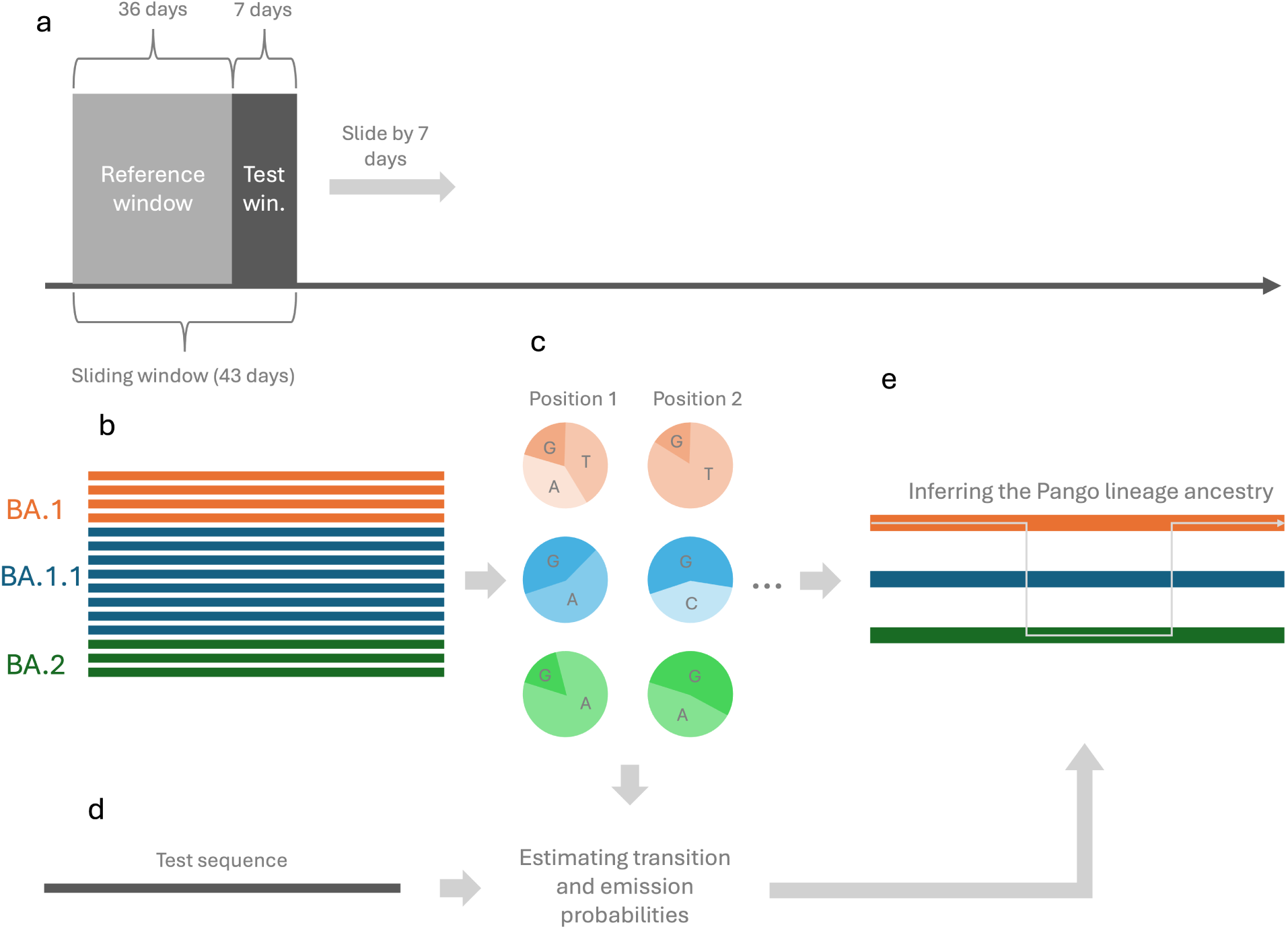
Overview of methods. (a) We first organize SARS-CoV-2 sequences collected in England between September 2020 and March 2024 into sliding windows of 43 days, which are advanced by 7 day increments. (b) In each sliding window, the first 36 days and last 7 days respectively comprise the reference window and test window. Sequences collected during the reference window comprise the reference set of sequences containing the mutational profile of each Pango lineage. (c) We next calculate the nucleotide frequency matrix containing per-position allele frequencies for each Pango lineage in the reference set. (d) For each test sequence collected during the test window, we use maximum likelihood to estimate frequencies that parameterize the transition and emission probabilities of our HMM. (e) We then use the Viterbi algorithm to predict the local Pango lineage ancestry for this sequence.

### 2.1 Obtaining SARS-CoV-2 sequences and clustering Pango lineages

We obtained SARS-CoV-2 sequences and metadata from GenBank, processed using the Nextstrain pipeline (Hadfield et al. 2018). After filtering for sequences collected in England between September 2020 and March 2024, we clustered Pango lineages based on their sequence count. We collapsed any Pango lineage with fewer than 10,000 sequences into its parental lineage, using unaliased Pango lineage names (O’Toole et al. 2021). This was done iteratively to ensure that all collapsed Pango lineages contained at least 10,000 sequences. Lineages without a defined parent were grouped into a shared “other” category. We collapsed 2,304 Pango lineages that existed during this period to 41 collapsed lineages (including the “other” category). Unless otherwise specified, all mentions of Pango lineages refer to the collapsed lineages resulting from this procedure.

### 2.2 Reference and test sets

From the sequences collected in England between September 2020 and March 2024, we generated sliding windows of reference and test set pairs. Each sliding window consisted of 43 days, and these windows were incremented by 7 days at a time to generate 185 sliding windows.

In each 43-day sliding window, the first 36 days and last 7 days respectively comprise the reference window and test window. Sequences collected during the reference and test windows respectively comprise the reference and test sets for this sliding window.

If more than 100,000 sequences were available during the reference window, we drew a random sample of 100,000 sequences and used this as the reference set. Similarly, if more than 3,000 sequences were available during the test window, we drew a random sample of 3,000 sequences and used this as the test set.

This process results in 185 pairs of reference and test sets. In the following sections, we describe our process for calculating the nucleotide frequency matrix for each reference set. We then define our HMM, which uses the nucleotide frequency matrix to predict the local Pango lineage ancestry for each sequence in the paired test set.

### 2.3 Calculating the nucleotide frequency matrix

For each of the 185 reference sets, we calculated a nucleotide frequency matrix that contains the frequency of each nucleotide (A, C, G, and T) at every genomic position for each Pango lineage. Nucleotide frequencies were calculated by dividing nucleotide counts at each position by the total sequence count within each Pango lineage. When calculating frequencies, we excluded all non-standard nucleotides (i.e., those other than A, C, G, and T). If no sequences in a Pango lineage carried any of the standard nucleotides at a position, we assigned equal probabilities (0.25 each) to A, C, G, and T.

### 2.4 Predicting the local Pango lineage ancestry

The local Pango lineage ancestry of a SARS-CoV-2 sequence refers to the sequence of Pango lineages ancestral to each genomic position of the SARS-CoV-2 sequence.

If a sequence derives from a recombination event between two sequences in two distinct Pango lineages, its local Pango lineage ancestry will consist of segments from these distinct lineages, with transitions between segments marking recombination breakpoints. Conversely, for non-recombinant sequences, the local Pango lineage ancestry will only contain a single parental lineage. It is important to note that the true local Pango lineage ancestry of a sequence in a test set is defined in relation to the Pango lineages present in the paired reference set. For example, lineages *L*_1_, *L*_2_, and *L*_3_ may all be present in the paired reference set, with lineage *L*_3_ arising from a recombination event between two sequences in lineages *L*_1_ and *L*_2_ respectively. Suppose that there is a sequence from lineage *L*_3_ in the test set. In this case, the true local Pango lineage ancestry of this sequence will have *L*_3_ as the lineage contributing ancestry at all genomic positions.

We predict the local Pango lineage ancestry of all sequences in each test set using the nucleotide frequency matrix calculated from the corresponding reference set and an HMM inspired by the Li and Stephens (2003) model.

### 2.5 Hidden Markov model to predict local Pango lineage ancestry

This HMM jointly models the latent local Pango lineage ancestry and the observed nucleotide sequence for each test sequence. It does so by considering three key components: (i) the probability of each lineage providing ancestry at the first position (initial state probabilities), (ii) the probability of transitioning between parental lineages from one position to the next (transition probabilities), and (iii) the probability of observing each nucleotide at a given position, conditional on the parental lineage (emission probabilities). Transitions between lineages correspond to recombination events.

Here, we define the HMM used to predict the local Pango lineage ancestry of a sequence in any given test set. We henceforth refer to this sequence as our test sequence. Let the genome length be denoted by *N* and let *t* ∈ {1, 2*, . . . , N* } index genomic positions. Our sequences are aligned, so all of our sequences have length *N* .

For our test sequence, we define the random variable for the parental Pango lineage at position *t* as *Z_t_*. *Z_t_* is supported on {1, 2*, . . . , M* }, where *M* is the number of distinct Pango lineages contained in the paired reference set (the reference set paired with the test set from which the test sequence is drawn). Each value of {1, 2*, . . . , M* } corresponds to one of these Pango lineages.

We further define, for the test sequence, the random variable for the observed nucleotide at position *t* as *O_t_*. *O_t_* is supported on {A, C, G, T}.

In the following sections, we define the three key components of this HMM, which are the initial state probabilities, the transition probabilities, and the emission probabilities.

#### 2.5.1 Initial state probabilities

The initial state probabilities give the probability of each Pango lineage being the parental lineage of the test sequence at the first genomic position. We define the initial state probability of Pango lineage *i* (*i* ∈ {1, 2*, . . . , M* }) as *π_i_* = *P* (*Z*_1_ = *i*). In our model, we set *π_i_* to the frequency of lineage *i* in the paired reference set. Let *n_i_* be the number of sequences assigned to lineage *i* in the reference set, and let 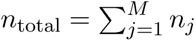 be the total number of sequences across all *M* lineages. Then,

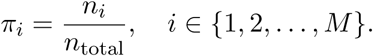

#### 2.5.2 Transition probabilities

The transition probabilities give the probability of transitioning from one parental Pango lineage to another between consecutive positions on the test sequence. Here, transitions between Pango lineages correspond to recombination breakpoints. We define the transition probability from Pango lineage *i* to Pango lineage *j* as

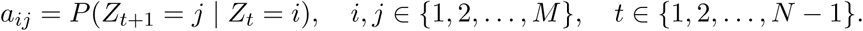

Here, *a_ij_* represents the probability that the parental Pango lineage of the test sequence changes from *i* to *j* between any consecutive positions on the genome. In our model, we set transition probabilities as

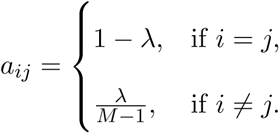

*λ* is the probability that there is a recombination breakpoint between consecutive positions on the genome. For the above formulation, we also assume that transitions between any two Pango lineages *i* ≠ *j* occur with the same probability. Because *λ* is an unknown parameter, we later describe our method for estimating *λ*.

#### 2.5.3 Emission probabilities

The emission probabilities give the probability of observing each nucleotide (i.e., A, C, G, or T) at a particular position on the test sequence, conditional on the parental Pango lineage at that position. We define the emission probability of observing nucleotide *k* at position *t*, conditional on the parental Pango lineage being *i* at position *t*, as

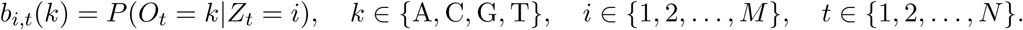

*b_i,t_*(*k*) depends on the nucleotide frequency matrix calculated from the paired reference set. We use *f_i,t_*(*k*) to denote the frequency of nucleotide *k* at position *t* in Pango lineage *i* in the paired reference set. To adjust for possible mutations and genotyping errors that could occur on the test sequence, we apply a pseudo-frequency *ϵ*. Specifically, we let

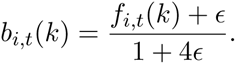

The pseudo-frequency *ϵ* assigns a non-zero probability of observing a nucleotide at position *t*, when the parental Pango lineage at *t* contains no sequences that have this nucleotide at *t* in the reference set. We want to allow for this non-zero probability in case the test sequence has acquired a mutation (or genotyping error) at position *t* that leads to an observed nucleotide that is not contained in the parental Pango lineage. A small value of *ϵ* allows occasional mutations or genotyping errors without forcing a lineage switch in the predicted local Pango lineage ancestry. Because *ϵ* is an unknown parameter, we describe our method for estimating *ϵ* in the following section.

We assume that positions with non-ACGT calls contain no information about the true nucleotide. Thus, we assign an emission probability of one to non-ACGT calls across all parental Pango lineages.

**Table 1.** Summary of symbols used in the HMM.

| Symbol | Description |
| --- | --- |
| $N$ | Genome length |
| $M$ | Number of Pango lineages in the reference set |
| $Z_t$ | Parental Pango lineage at position $t$ |
| $O_t$ | Observed nucleotide at position $t$ |
| $\lambda$ | Transition probability |
| $\epsilon$ | Pseudo-frequency for emissions (accounts for mutations and genotyping errors) |
| $a_{ij}$ | Transition probability from lineage $i$ to lineage $j$ |
| $b_{i,t}(k)$ | Emission probability of nucleotide $k$ at $t$ given $Z_t = i$ |
| $f_{i,t}(k)$ | Frequency of nucleotide $k$ at $t$ in lineage $i$ in the reference set |
| $\pi_i$ | Initial state probability that $Z_1 = i$ |

### 2.6 Maximum likelihood estimation of parameters in the hidden Markov model

We have two unknown parameters in our HMM. *λ* is the probability that the parental Pango lineage changes between consecutive positions and *ϵ* is our pseudo-frequency, which adjusts emission probabilities to accommodate mutations or genotyping errors on the test sequence.

To perform maximum likelihood estimation on these two parameters, we first obtain the probability of the observed nucleotide sequence of the test sequence, conditional on these two parameters. In this section, we describe the procedure we use to obtain this probability.

Using the transition and emission probabilities described in the previous sections, it is relatively straightforward to obtain the joint probability of a candidate local Pango lineage ancestry and the observed nucleotide sequence for a test sequence. Let *i*_1:*N*_ = (*i*_1_*, i*_2_*, . . . , i_N_* ) ∈ [*M* ]*^N^* be a candidate local Pango lineage ancestry and *k*_1:*N*_ = (*k*_1_*, k*_2_*, . . . , k_N_* ) ∈ {A, C, G, T}*^N^* be the observed nucleotide sequence of this test sequence. Finally, let *Z*_1:*N*_ = (*Z*_1_*, Z*_2_*, . . . , Z_N_* ) and *O*_1:*N*_ = (*O*_1_*, O*_2_*, . . . , O_N_* ). Then,

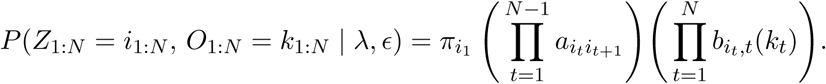

To obtain the marginal probability of the observed nucleotide sequence, we can simply sum up this joint probability across all possible local Pango lineage ancestries, as shown below.

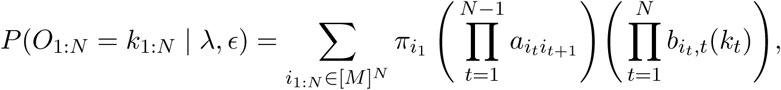

This procedure can be carried out efficiently using the forward algorithm described in Rabiner (1989). We implemented a fast version of this forward algorithm that computes the induction step in O(*M* ) time compared to the normal O(*M* ^2^) time (see Section S1 of the Supplementary Materials). We can maximize this marginal probability with respect to our two parameters to obtain our maximum likelihood estimates, as shown below.

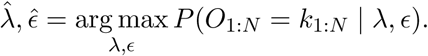

Maximum likelihood estimation of *λ* and *ϵ* is done for each test sequence. Optimization was carried out with the limited-memory BFGS algorithm subject to box constraints, using scipy.optimize.minimize (method = "L-BFGS-B") (Virtanen et al. 2020). Our version of the forward algorithm results in a large reduction in computation time, because the marginal likelihood is evaluated repeatedly during L-BFGS-B optimization.

During numerical optimization, we reparameterize *λ* to *τ* = *λ*(*N* −1), which represents the expected number of transitions for the test sequence. Furthermore, we optimized *ϵ* on the log scale and later exponentiated to obtain our estimate in the original scale. The search was initialized at (log(*ϵ*)*, τ* ) = (log(0.005), 1) and restricted to the intervals log(*ɛ*) ∈ [log(10^−8^), log(0.02)] and *τ* ∈ [0, 3]. The reparameterization of *λ* to *τ* was done to avoid possible numerical instabilities that might arise when trying to optimize *λ* directly, because we expect *λ* to be close to zero. We similarly optimized *ɛ* in the log scale because we expect *ϵ* to be close to zero.

We chose the upper bound of three for *τ* because most discovered recombinant lineages were detected to have three or fewer breakpoints. However, this does not prevent the predicted local Pango lineage ancestry from having more than three breakpoints.

### 2.7 Obtaining the most likely sequence of Pango lineage ancestry

We apply the Viterbi algorithm described in Rabiner (1989) to each test sequence to obtain the most probable sequence of parental Pango lineage states along the genome,

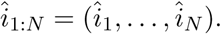

This represents the predicted local Pango lineage ancestry for this test sequence. Transitions between Pango lineage ancestries represent predicted recombination breakpoints.

When applying the Viterbi algorithm, we use our maximum likelihood estimates of the two frequencies, *λ* and *ϵ*, described in earlier sections. Specifically, we compute the sequence of Pango lineage ancestry that maximizes the joint probability of the ancestry path and the observed nucleotide sequence, given our maximum likelihood estimates. In other words,

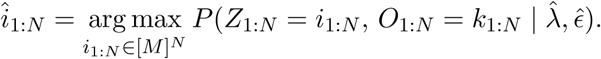

Note that because *P* (*O*_1:*N*_ = *k*_1:*N*_ | *λ*^^^*, ɛ*^) does not depend on the local Pango lineage ancestry, the above is equivalent to maximizing the posterior probability of the local Pango lineage ancestry given the observed nucleotide sequence and our maximum likelihood estimates. In other words,

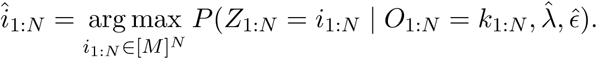

### 2.8 Simulation study

To assess our method’s ability to detect recombination and accurately predict local Pango lineage ancestries, we conducted a simulation study using synthetic SARS-CoV-2 sequences with known local Pango lineage ancestries. These synthetic sequences were generated from real SARS-CoV-2 genomes.

We generated these synthetic sequences using the reference set comprised of 14,599 SARS-CoV-2 sequences collected in England between November 6, 2022 and December 11, 2022. We simulated 1,000 recombinant sequences with two parental lineages and 1,000 control sequences with one parental lineage.

We generated 500 recombinant sequences using a single recombination breakpoint. To generate these sequences, we randomly sampled two parental sequences from two different Pango lineages in the reference set and copied nucleotides from one parent up to a breakpoint randomly chosen on the genome, and from the other parent thereafter. We generated the remaining 500 recombinant sequences using two breakpoints. For these sequences, we again sampled two parental sequences from different Pango lineages. We chose two breakpoints randomly from all possible breakpoint combinations on the genome and inserted a middle segment from one sequence between these breakpoints, replacing the corresponding region in the genome of the other sequence. If a synthetic recombinant sequence was identical to or differed by only one mutation from one of its parental sequences, we discarded this sequence and repeated the sequence generation process. We obtained 1,000 control sequences by sampling 1,000 sequences at random from the reference set.

To mimic mutations on all 2,000 synthetic sequences, we drew the mutation count from an empirical distribution obtained by tallying nucleotide substitution counts on each branch of a tree containing one tip per Pango lineage (Hadfield et al. 2018). The empirical distribution of nucleotide substitution counts was right-skewed (n = 567; median = 2 [IQR 1–4]; mean = 3.45; 95th percentile = 8; range 1–70). Given the drawn mutation count *m*, we sampled *m* genomic positions uniformly at random and replaced the existing nucleotide at each position with one of the other four nucleotides (A, C, G, T, or N, excluding the original base) chosen at random. In our aligned sequences, N represents an unknown nucleotide.

For each of the 2,000 synthetic sequences, we applied the method described in Section 2.6 to estimate the transition probability *λ* and pseudo-frequency *ϵ*. We then predicted the local Pango lineage ancestry for each sequence using the method described in Section 2.7. Emission probabilities for the HMM were based on the nucleotide frequency matrix calculated from the reference set of sequences collected between November 6, 2022 and December 11, 2022.

To evaluate performance, we conducted several quantitative assessments. First, we estimated the sensitivity and specificity of our method for classifying sequences as recombinant or non-recombinant. We classified a synthetic sequence as a recombinant if the predicted local Pango lineage ancestry contained at least one lineage transition.

Second, we calculated the mean position-by-position accuracy of the predicted local Pango lineage ancestry across synthetic sequences by comparing the predicted parental lineage at each genomic position to the true parental lineage.

Third, we assessed whether the correct parental lineage (for control sequences) or lineage pair (for recombinant sequences) was correctly recovered. We calculated the proportion of synthetic sequences for which the parental lineage or lineage pair was correctly recovered. Both the mean position-by-position accuracy and the recovery rate of parental lineages were calculated separately for recombinant and non-recombinant sequences.

For the sensitivity, specificity, and recovery rate of parental Pango lineages, we report 95% exact binomial confidence intervals. For mean position-by-position accuracy, we calculated 95% bootstrap confidence intervals by sampling 500 times with replacement from synthetic sequences (either from the set of recombinants or the set of non-recombinant sequences), calculating the mean position-by-position accuracy in each bootstrap sample, and taking the 2.5th and 97.5th percentiles of the bootstrapped estimates.

We next quantified the association between the sensitivity to detect recombinants *s*(*d*) and the genome-wide Hamming distance *d* between the parental sequences of recombinants using a logistic regression, which can be written as,

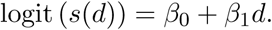

It follows that exp(*β*_1_) represents the multiplicative difference in the odds of detection for two synthetic recombinant sequences whose parental Hamming distances are one unit apart. To fit this logistic regression, we used the HMM’s predicted label for all 1,000 synthetic recombinants (1 if the model detected the sequence as a recombinant and 0 otherwise) as the outcome.

To quantify the accuracy of predicted breakpoint positions, we calculated the distance between each predicted breakpoint position and its corresponding true genomic position. We restricted analysis to synthetic recombinants whose detected breakpoint count matched the number of true breakpoints. For synthetic recombinants with one true breakpoint, we calculated the distance between the true and detected breakpoint position. For synthetic recombinants with two true breakpoints, we ordered true and detected breakpoint positions 5’ to 3’, paired them positionally (first with first, second with second), and calculated the distance between each pair. We then calculated the mean breakpoint distance separately for recombinants with one and two breakpoints.

We obtained 95% confidence intervals for the mean breakpoint distance via nonparametric bootstrap. Specifically, we sampled recombinant sequences with replacement 500 times within each stratum (one and two breakpoints), calculated the mean breakpoint distance for each bootstrap sample, and took the 2.5th and 97.5th percentiles of the bootstrapped estimates within each stratum. When detected and true breakpoints counts differed, we did not compute a distance. Instead we recorded the detected and true breakpoint counts per sequence and summarized mismatches in a contingency table.

### 2.9 Empirical data analysis

We applied our method to the full set of SARS-CoV-2 sequences collected in England between September 2020 and March 2024. We describe how we obtain these sequences in Section 2.1. As described in Section 2.2, sequences were divided into temporally matched reference and test sets using a 43-day sliding window. For each sequence in each test set, we estimated the transition probability *λ* and pseudo-frequency *ϵ* using maximum likelihood (see Section 2.6). We then predicted the local Pango lineage ancestry for each sequence using the Viterbi algorithm (see Section 2.7). There were 440,307 sequences across all test sets.

We classified any sequence with one or more lineage transitions in their predicted local Pango lineage ancestry as recombinant. For each 7-day test window, we calculated the detected recombinant proportion, or the number of detected recombinants divided by the total number of tested sequences from this window. Recall that the total number of tested sequences can vary across test windows (see Section 2.2).

We hypothesized that the detected recombinant proportion would be positively associated with community SARS-CoV-2 prevalence across test windows. This is because co-infection by two distinct Pango lineages, which is required for the emergence of detectable recombinant sequences, occurs more frequently when community prevalence of SARS-CoV-2 is high. To evaluate our hypothesis, we compared the detected recombinant proportion in each test window with community prevalence estimated from the UK Office for National Statistics (ONS) Coronavirus Infection Survey (Pouwels et al. 2021). ONS provides prevalence estimates by date. For comparability, we computed the mean ONS prevalence estimate within each test window and used this window-averaged prevalence estimate in our analysis.

It is important to note that our method does not detect recombinants whose parental sequences belong to the same Pango lineage. Thus, the detected recombinant proportion in a given test window reflects only recombination events occurring between distinct Pango lineages. Because overall SARS-CoV-2 community prevalence does not consider the relative frequencies of circulating lineages, we do not expect the detected recombinant proportion to be strongly associated with unadjusted prevalence estimates. Hypothetically, if only one lineage is circulating, we would not detect any recombination even if community prevalence of SARS-CoV-2 is high.

To address this, we also consider a lineage-adjusted measure of prevalence that incorporates the joint circulation of distinct Pango lineages, defined in equation S3 of the Supplementary Materials. As shown in Section S2 of the Supplementary Materials, this quantity corresponds to an estimate of the expected true positive recombinant proportion in each test window under a model of independent co-infection (Chin et al. 2024). Comparing this quantity with the detected recombinant proportion allows us to assess how the observed detection frequency of recombinants in each test window relates to the expected opportunity for lineage co-infection within that window. For this comparison, because the expected true positive recombinant proportion is based on pairwise lineage co-circulation, we recalculated the detected recombinant proportion as the number of detected recombinants with two inferred parental lineages divided by the total number of tested sequences in each window.

We next counted the number of detected recombinant sequences across all test windows, stratified by unique parental Pango lineage pairs (e.g., BA.1.1–BA.2). For each lineage pair, we compared the observed number of detected recombinants to an estimated number of true positive recombinants derived from the same independent co-infection model as before. Specifically, we used observed lineage proportions and ONS prevalence estimates to compute the expected number of co-infections involving a given pair of distinct lineages in each test window, and aggregated these expectations across windows to obtain a predicted true positive count for each parental lineage pair (see Section S2 and equation S4 of the Supplementary Materials). This comparison allows us to assess how the observed detection count of recombinants with each lineage pair relates to its expected opportunity for co-infection.

Finally, we aggregated inferred breakpoint positions across all detected recombinant sequences to obtain the empirical genome-wide distribution of recombination breakpoints. We examined this distribution to identify genomic regions in which recombination breakpoints were enriched.

## 3 Results

### 3.1 Simulation study

To evaluate the performance of our method for detecting recombinant SARS-CoV-2 sequences, we conducted a series of assessments based on the predicted local Pango lineage ancestry of synthetic SARS-CoV-2 sequences. The process used to generate synthetic sequences are described in Section 2.8.

First, we estimated the sensitivity and specificity of our method for classifying a sequence as a recombinant or non-recombinant. We classified a test sequence as a recombinant if the predicted local Pango lineage ancestry contained at least one lineage transition. Our method achieved a sensitivity of 0.801 (95% CI: [0.775, 0.825]) and a specificity of 0.989 (95% CI: [0.980, 0.994]).

To assess the accuracy of predicted local Pango lineage ancestries, we computed the mean position-by-position accuracy separately for recombinant and control sequences. On average, the inferred Pango lineage matched the true parental lineage at 86.9% (95% CI: [85.9%, 87.9%]) of genomic positions for recombinant sequences. Among control sequences, mean position-by-position accuracy was 99.2% (95% CI: [98.6%, 99.7%]).

We further evaluated how often the true parental lineage pair or lineage was recovered for recombinant and control sequences respectively. In 69.9% (95% CI: [67.0%, 72.7%]) of synthetic recombinant sequences, we detected two parental lineages that matched the true parental lineage pair. There was an overlap between the true and detected lineages in 100% (95% CI: [99.6%, 100%]) of synthetic recombinant sequences. In 98.4% (95% CI: [97.4%, 99.1%]) of synthetic control sequences, we detected a single parental lineage that matched the true parental lineage, and there was an overlap between the true and detected lineages in 99.4% (95% CI: [98.7%, 99.8%]) of synthetic control sequences.

In Table 2, we report how often the true parental lineage pair was recovered for recombinant sequences, stratifying by the true parental lineage pair. We counted the number of times we detected two parental lineages that matched the true parental lineage pair, for recombinants with each true parental lineage pair. We restricted this analysis to true parental lineage pairs with at least ten synthetic recombinants.

**Table 2.** Detection of parental lineages for recombinant sequences, stratified by true parental lineage pair. We report 95% exact binomial confidence intervals for the proportion of sequences for which we detected two parental lineages that matched the true parental lineage pair.

| True lineages | Num. samples | Recovered | Prop. | 2.5 % CI | 97.5 % CI |
| --- | --- | --- | --- | --- | --- |
| (BA.5.2, BQ.1.1) | 81 | 76 | 0.938 | 0.862 | 0.980 |
| (BA.5.2, CH.1.1) | 13 | 12 | 0.923 | 0.640 | 0.998 |
| (BQ.1.1, CH.1.1) | 72 | 65 | 0.903 | 0.810 | 0.960 |
| (BA.4, BQ.1.1) | 18 | 16 | 0.889 | 0.653 | 0.986 |
| (BA.2, BQ.1.1) | 89 | 78 | 0.876 | 0.790 | 0.937 |
| (BA.5.1, BQ.1.1) | 16 | 14 | 0.875 | 0.617 | 0.984 |
| (BQ.1.1, XBB.1) | 31 | 27 | 0.871 | 0.702 | 0.964 |
| (BA.5, BE.1.1) | 11 | 9 | 0.818 | 0.482 | 0.977 |
| (BA.5.2.1, BQ.1.1) | 78 | 63 | 0.808 | 0.703 | 0.888 |
| (BA.2, BA.5.2.1) | 27 | 21 | 0.778 | 0.577 | 0.914 |
| (BA.5.2.1, CH.1.1) | 13 | 10 | 0.769 | 0.462 | 0.950 |
| (BA.2, BA.5.2) | 21 | 16 | 0.762 | 0.528 | 0.918 |
| (BA.5.2, BE.1.1) | 40 | 30 | 0.750 | 0.588 | 0.873 |
| (BE.1.1, XBB.1) | 11 | 8 | 0.727 | 0.390 | 0.940 |
| (BA.2, BE.1.1) | 46 | 33 | 0.717 | 0.565 | 0.840 |
| (BA.2, XBB.1) | 10 | 7 | 0.700 | 0.348 | 0.933 |
| (BA.5.1, BE.1.1) | 13 | 9 | 0.692 | 0.386 | 0.909 |
| (BE.1.1, CH.1.1) | 32 | 22 | 0.688 | 0.500 | 0.839 |
| (BA.5.2.1, BE.1.1) | 30 | 20 | 0.667 | 0.472 | 0.827 |
| (BA.2, CH.1.1) | 14 | 9 | 0.643 | 0.351 | 0.872 |
| (BA.5, BQ.1.1) | 22 | 12 | 0.545 | 0.322 | 0.756 |
| (BA.5.2, BA.5.2.1) | 19 | 9 | 0.474 | 0.244 | 0.711 |
| (BQ.1.1, other) | 23 | 10 | 0.435 | 0.232 | 0.655 |
| (BE.1.1, BQ.1.1) | 128 | 24 | 0.188 | 0.124 | 0.266 |

We then evaluated whether the sensitivity to detect synthetic recombinant sequences was associated with the Hamming distance between the two parental sequences of each synthetic recombinant sequence. Using logistic regression, we found a positive association between the parental Hamming distance and the sensitivity (*p <* 2 × 10^−16^ using a two-sided Wald test). We estimate that for two recombinant sequences that differ by one unit in their parental Hamming distances, the odds of detection is 1.11 times higher in the recombinant sequence with the higher parental Hamming distance (95% CI: [1.09, 1.13]). The relationship between the parental Hamming distance and detection probability is shown in Figure 2, which displays the fitted logistic regression and the associated 95% pointwise confidence band.

**Figure 2.**
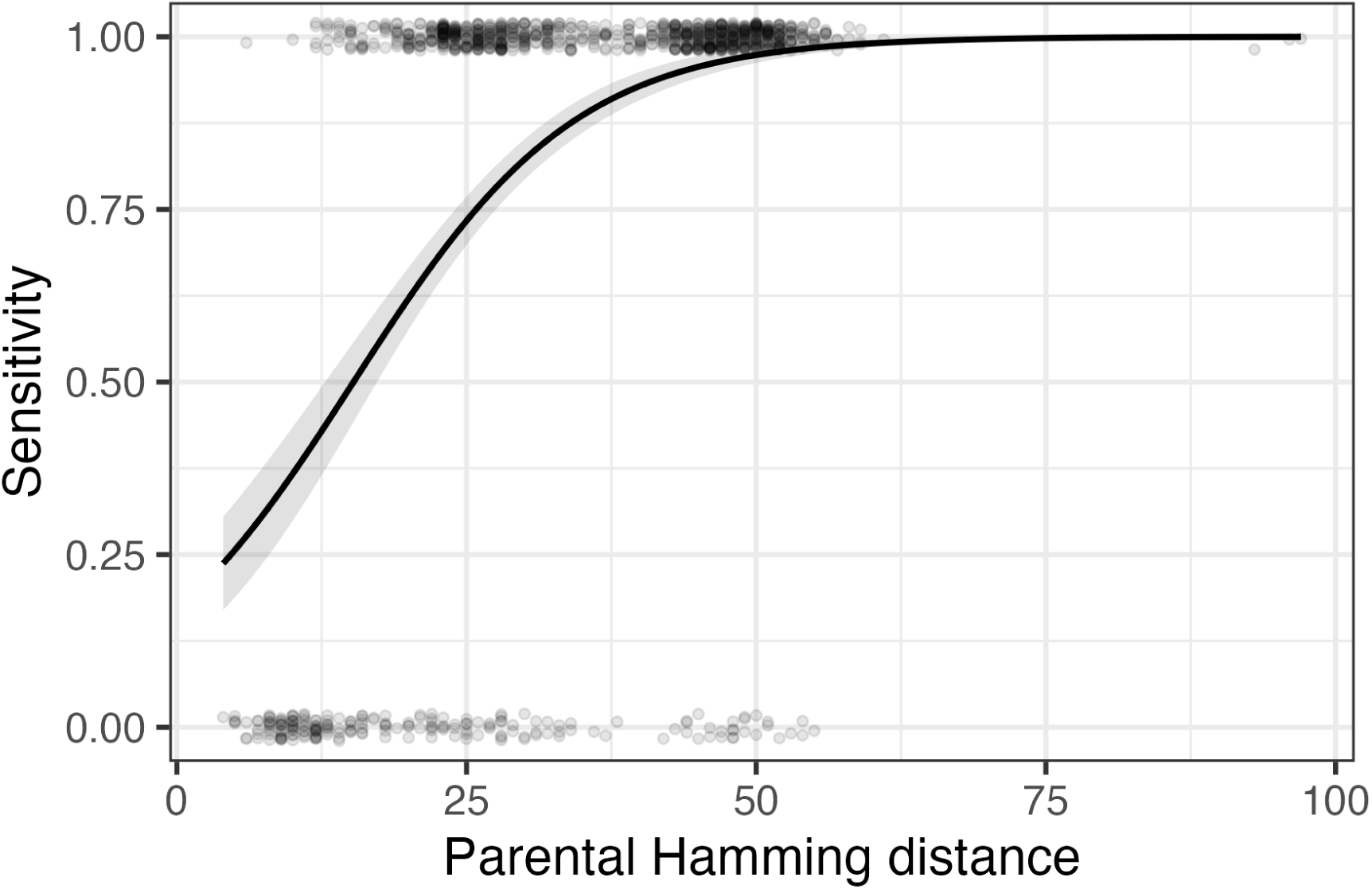
Sensitivity to detect recombinants as a function of the Hamming distance between their two parental sequences. Each point represents a synthetic recombinant sequence, with *y* = 1 indicating that it was classified as a recombinant using our method. *y* values are jittered vertically to avoid overplotting. The black line represents the logistic regression fit. The grey band represents pointwise 95% CIs.

We next assessed the accuracy of predicted breakpoint positions. Among sequences with one break-point (that had one predicted breakpoint), the mean breakpoint distance was 1238 nucleotides (95% CI: [1108, 1386]). For sequences with two breakpoints (that had two predicted breakpoints), the mean breakpoint distance was 1007 nucleotides (95% CI: [901, 1125]). For synthetic recombinants with two true breakpoints, we ordered true and detected breakpoint positions 5’ to 3’, paired them positionally (first with first, second with second), and calculated the distance between each pair. We then averaged the paired breakpoint distances across all sequences.

It is difficult to assess the accuracy of predicted breakpoint positions for a recombinant sequence whose predicted breakpoint count does not match its true breakpoint count. A confusion matrix of predicted and true breakpoint counts for recombinant sequences is shown in Table 3.

**Table 3.**
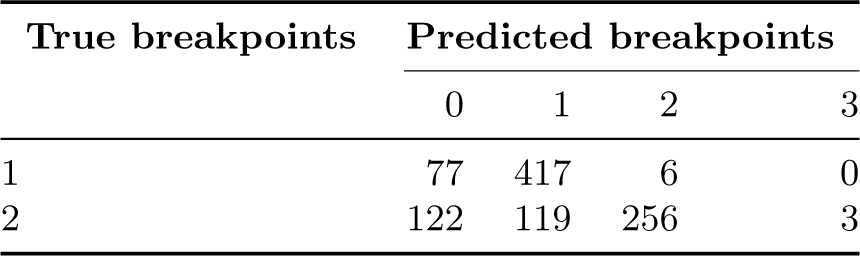
Confusion matrix of predicted versus true breakpoint counts.

### 3.2 Empirical data analysis

We used our method to predict the local Pango lineage ancestry for 440,307 SARS-CoV-2 sequences collected in England between September 2020 and March 2024. These sequences were sampled across 185 test windows that each consisted of a 7-day period with no gaps between successive windows. Of the 440,307 sequences, 7619 were detected to be recombinant sequences using our method, which corresponds to 1.73% (95% CI: [1.69%, 1.77%]) of sequences.

In Figure 3, we plot the detected recombinant proportion (the proportion of tested sequences detected to be recombinant using our method) in each test window and SARS-CoV-2 prevalence estimates from the UK Office for National Statistics (ONS) Coronavirus Infection Survey (Pouwels et al. 2021), averaged within each test window. We see a positive trend in the detected recombinant proportion over time. We observe weak positive correlation between the detected recombinant proportion and ONS prevalence estimates (Pearson correlation *r* = 0.18). Because ONS prevalence estimates were only available from June 2020 to March 2023, the Pearson correlation was calculated using the 132 test windows in which ONS prevalence estimates were available. Fitting a linear regression of the detected recombinant proportion on the ONS prevalence (Figure 3, bottom panel), we observed a positive association between the two quantities (*β* = 0.10; HC3 heteroskedasticity-robust *p* = 0.0377; *N* = 132).

**Figure 3.**
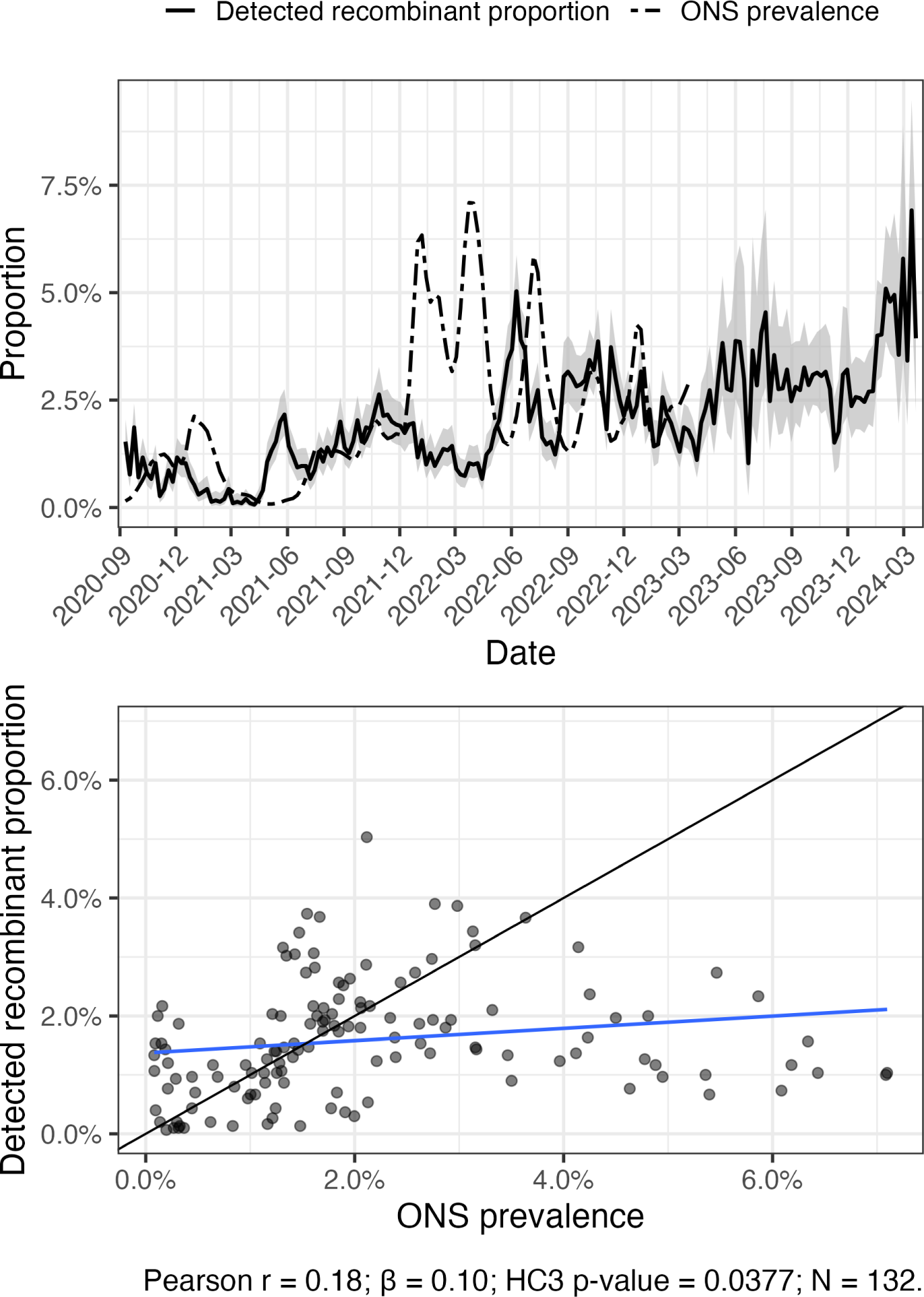
Comparison of detected recombinant proportion and ONS prevalence estimates across test windows. (Top) Time series showing the detected recombinant proportion (solid line), with shaded bands indicating 95% exact binomial confidence intervals. The dashed line shows ONS prevalence estimates. (Bottom) Scatterplot comparing the detected recombinant proportion with ONS prevalence estimates across test windows. The blue line shows the fitted linear regression. The black line is the identity line.

As noted in Section 2.9, overall community prevalence does not account for the joint circulation of distinct parental lineages. Thus, we do not necessarily expect overall community prevalence to be strongly correlated with the detected recombinant proportion, which only reflects recombination events between distinct Pango lineages because recombinants whose parental sequences belong to the same lineage cannot be detected by our method. We next introduce a lineage-adjusted prevalence measure, *ρ*^(*w*), that explicitly incorporates pairwise Pango lineage frequencies (see equation S3 of the Supplementary Materials). As shown in Section S2 of the Supplementary Materials, this quantity estimates the expected proportion of true positive recombinants in each test window.

Because the expected proportion of true positive recombinants is based on pairwise lineage cocirculation, we recalculate the detected recombinant proportion in each window as the number of detected recombinants with two inferred parental lineages divided by the total number of tested sequences. For reference, out of the 7619 sequences detected to be recombinant across all test windows, 7063 had two inferred parental lineages, corresponding to 92.7% (95% CI: [92.1%, 93.3%]) of sequences. In Figure 4, we compare this detected recombinant proportion with the estimated expected proportion of true positive recombinants across test windows.

**Figure 4.**
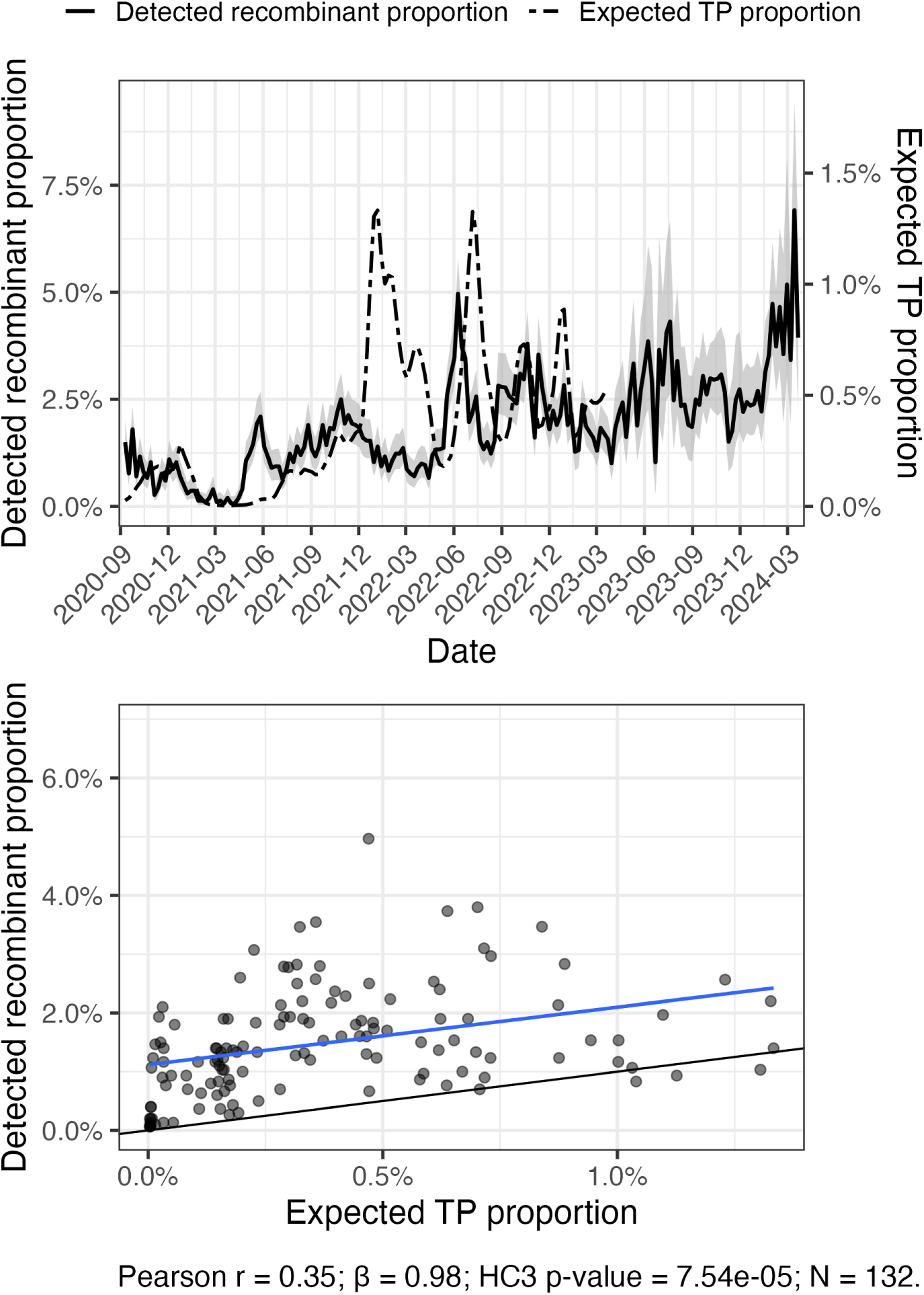
Comparison of detected recombinant proportion and estimated expected true positive recombinant proportion across test windows. (Top) Time series showing the detected recombinant proportion in each test window (solid line), with shaded bands indicating 95% exact binomial confidence intervals; values are shown on the left axis. The dashed line shows the estimated expected true positive recombinant proportion, with values shown on the right axis. (Bottom) Scatterplot comparing the detected recombinant proportion with the estimated expected true positive recombinant proportion across test windows. The blue line shows the fitted linear regression. The black line is the identity line.

Fitting a linear regression of the detected recombinant proportion on the estimated expected true positive recombinant proportion (Figure 4, bottom panel), we observed a strong positive association between the two quantities (*β* = 0.98; HC3 heteroskedasticity-robust *p* = 7.54 × 10^−5^; *N* = 132).

This suggests that the detected recombinant proportion is strongly associated with expectations based on co-infection opportunities between distinct Pango lineages.

The detected recombinant proportion almost always exceeds the estimated expected true positive recombinant proportion within individual test windows, as evidenced by most points falling above the identity line in the bottom panel of Figure 4. This is expected because the former proportion includes both true positives and false positives detected by our method.

We next counted the number of detected recombinants stratified by unique parental lineage pairs.

For each parental lineage pair *i < j*, we compared the number of detected *i*–*j* recombinants (excluding cases in which one of the parental lineages was in the “other” category) to E^^^[*R_i,j_*] derived in equation S4 of the Supplementary Materials, which represents our estimate of the expected number of true positive *i*–*j* recombinants detected across all test windows. For brevity, we henceforth refer to E^^^[*R_i,j_*] as the expected true positive count for *i*–*j* recombinants.

ONS SARS-CoV-2 prevalence estimates are used to obtain expected true positive counts for each lineage pair *i < j*. For comparability with these counts, detected recombinant counts are also restricted to windows with available ONS prevalence estimates (until the window ending March 19, 2023). For reference, test windows before March 19, 2023 contained 387,054 sequences, with 5006 sequences detected as recombinant with exactly two parental lineages in their predicted local Pango lineage ancestry (excluding cases in which one of the parental lineages were in the “other” category).

In Figure 5, we plot the observed number of detected *i*–*j* recombinants against the corresponding expected true positive counts E^^^[*R_i,j_*] across parental lineage pairs. Fitting a linear regression of detected recombinant counts on expected true positive counts across lineage pairs (*i < j*), we found a strong positive association (*β* = 1.79; HC3 heteroskedasticity-robust *p* = 2.04 × 10^−6^; *N* = 279).

**Figure 5.**
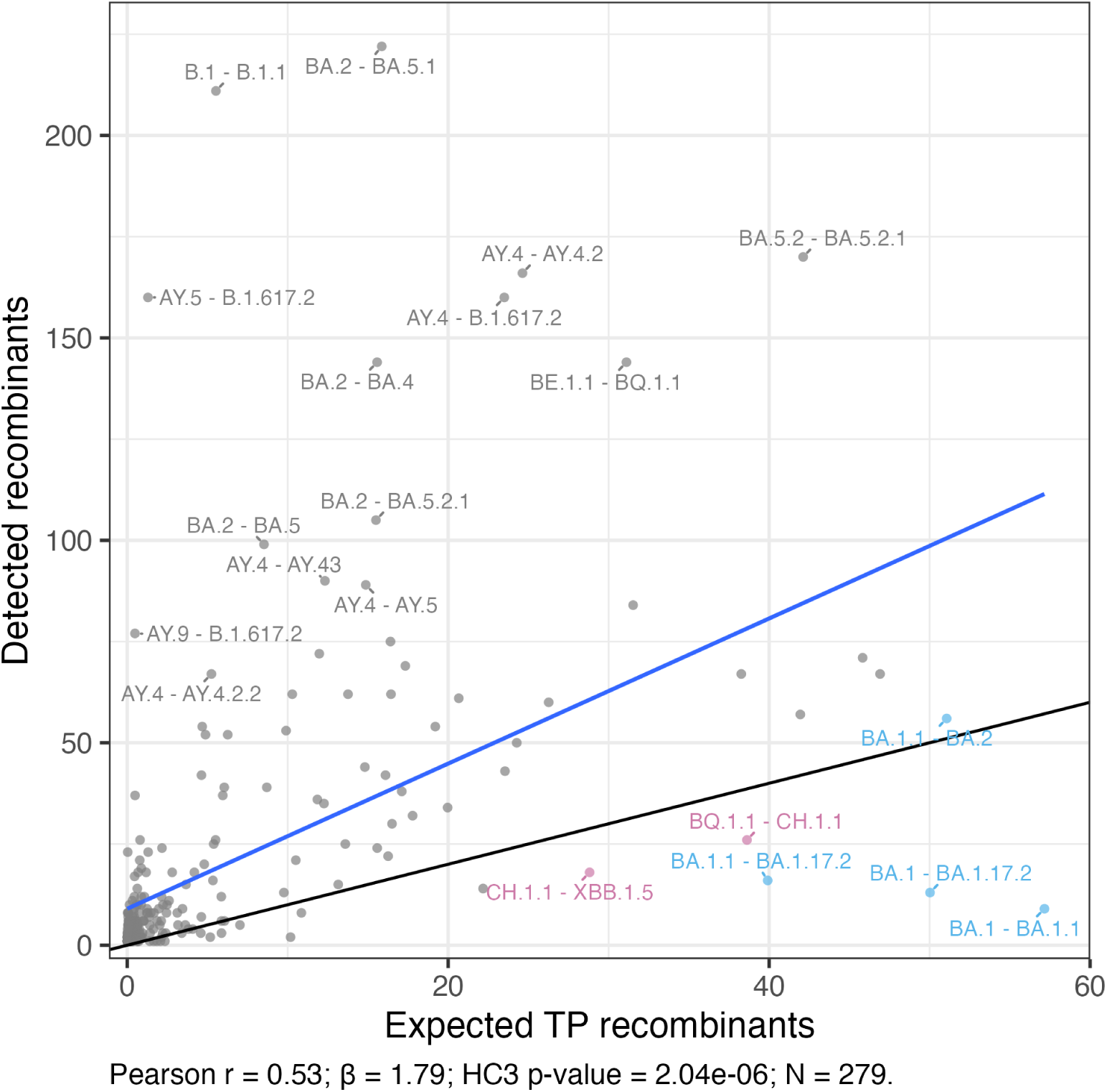
Scatterplot of detected recombinant counts against expected true positive counts (Ê[*R_i,j_*]). Each point represents a parental lineage pair. The blue line shows the ordinary least squares fit. The black line represents the identity line. We label the 20 parental lineage pairs with the largest absolute ordinary least squares residuals. Under-represented lineage pairs are colored. Light blue indicates parental lineage pairs that co-occurred during late 2021 to early 2022, whereas purple indicates pairs that co-occurred during late 2022 to early 2023.

Because expected true positive counts do not include contributions from false positives, we generally expect the number of detected recombinants to be larger across parental lineage pairs *i < j*. We observe this in Figure 5 (most points fall above the identity line).

Several parental lineage pairs deviate markedly from the overall relationship between detected counts and expected true positive counts. We identified the 20 parental lineage pairs with the highest absolute residual values and annotated them in Figure 5.

Six of these parental lineage pairs had negative residual values. We refer to these parental lineage pairs as under-represented lineage pairs. Recombinants with these parental lineage pairs (except for BA.1.1–BA.2) had lower detected counts than expected true positive counts (these recombinants fall below the identity line in Figure 5). This should be interpreted with caution. This apparent under-representation may be attributable to uncertainty in expected true positive counts or sampling variability in detected counts. Alternatively, this may suggest recombinants with these parental lineage pairs occur less frequently in the population than expected under our co-infection model or we have lower sensitivity to detect these recombinants compared to our overall sensitivity (see Section S2 of the Supplementary Materials).

We likely have low sensitivity to detect BA.1–BA.1.1, BA.1–BA.1.17.2, and BA.1.1–BA.1.17.2 recombinants, given that only a few mutations separate these lineage pairs. Using consensus sequences and ignoring non-standard nucleotides, pairwise Hamming distances were one for BA.1–BA.1.1, three for BA.1–BA.1.17.2, and four for BA.1.1–BA.1.17.2. Recall that in our simulation study, the sensitivity of our method was low when parental sequences only differed by a few mutations (see Figure 2).

Pairwise Hamming distances were relatively high for BQ.1.1–CH.1.1 (39 nucleotides), BA.1.1–BA.2 (41 nucleotides), and CH.1.1–XBB.1.5 (34 nucleotides). However, the sensitivity to detect recombinants with these parental lineage pairs may still be low, depending on where recombinantion breakpoints occur. The resulting recombinant sequence may only have a few mutations relative to one of its parents, if most of its genome is inherited from this parent. Furthermore, these parental lineage pairs lie close to the identity line *y* = *x* (slightly above the line for BA.1.1–BA.2). Thus, their under-representation may be explained by uncertainty in expected true positive counts or sampling variability in detected counts.

Interestingly, every under-represented parental lineage pair co-circulated in England during two intervals, late 2021 to early 2022 (BA.1–BA.1.1, BA.1–BA.1.17.2, BA.1.1–BA.1.17.2, BA.1.1–BA.2) and late 2022 to early 2023 (BQ.1.1–CH.1.1, CH.1.1–XBB.1.5), when the detected recombinant proportion was low relative to ONS prevalence estimates (see Figure 3, top panel). For under-represented parental lineage pairs, Figure 6 shows the pairwise product of their lineage frequencies across test windows. Recombinants with these parental lineage pairs had few detected counts during these two periods, indicating that these under-represented lineage pairs contributed to the lower frequency of recombinants detected during this period. In particular, during late 2021 to early 2022, it is likely that the co-circulation of closely related Omicron sublineages, specifically BA.1– BA.1.1, BA.1–BA.1.17.2, and BA.1.1–BA.1.17.2, contributed to the small detected proportion of recombinants during this period, relative to the overall relationship between the detected proportion and expected true positive proportion (see Figure 4, top panel). This is consistent with lower sensitivity to detect recombinants arising from parental lineages with high sequence similarity.

**Figure 6.**
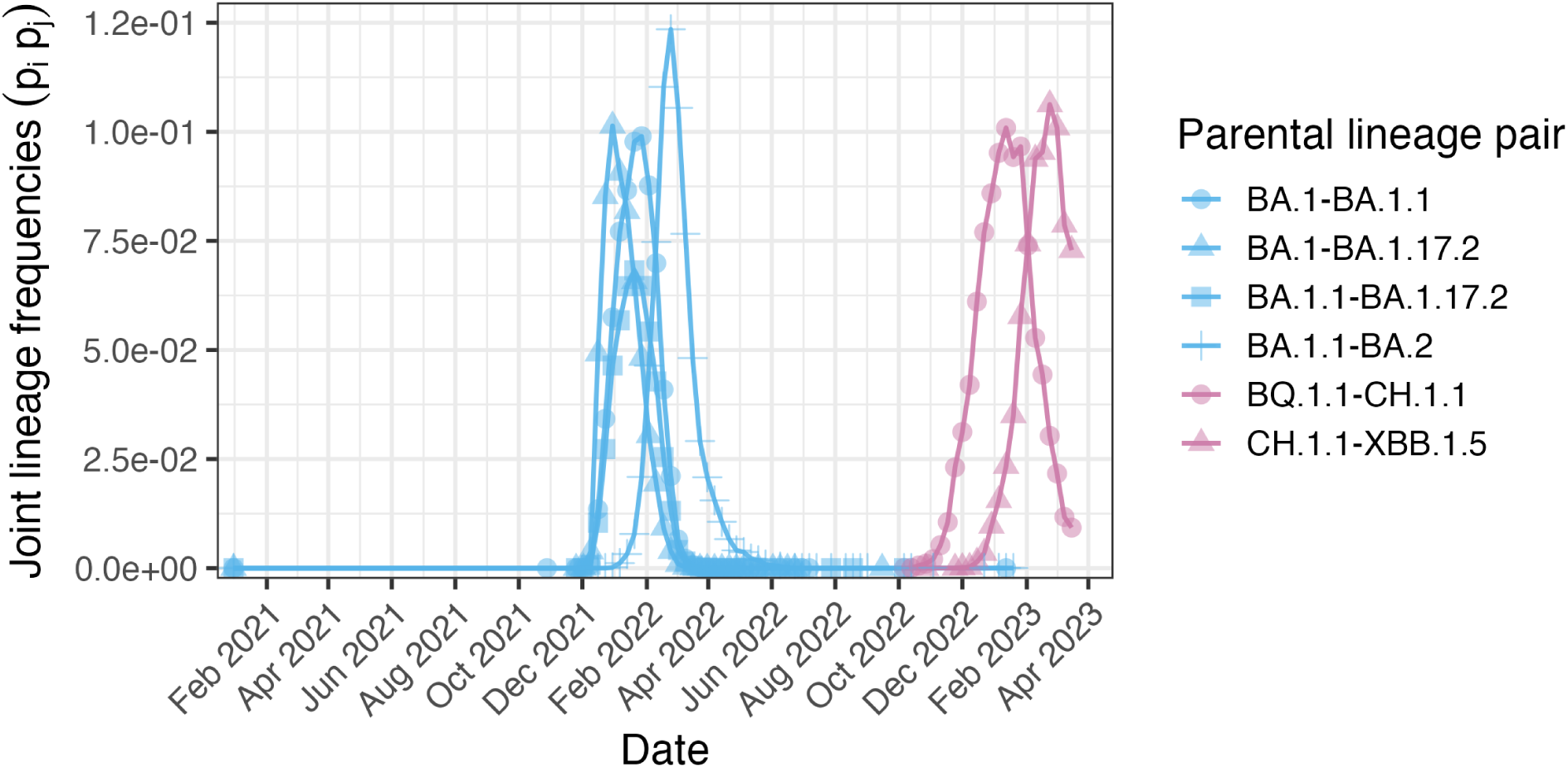
Joint frequencies across test windows for under-represented lineage pairs.

In the process of estimating the expected true positive recombinant proportion, we estimated the detection factor *θ* and false positive rate *ϕ* to be 0.557 (95% CI: [0.297, 0.817]) and 0.011 (95% CI: [0.009, 0.013]) respectively. Recall that *θ* equals sensitivity times the probability that a sample from a co-infected individual is a recombinant (see Section S2 of the Supplementary Materials). Confidence intervals are Wald intervals from the linear regression model described by equation S2 of the Supplementary Materials treating *x_w_* as fixed and using HC3 standard errors. This linear regression is shown in Figure S1 of the Supplementary Materials. Our estimated false positive rate closely matches what we estimated in the simulation (see Section 3.1).

Across 7619 detected recombinants, we inferred 9105 recombination breakpoints. 6324 detected recombinants (83.0%) had one breakpoint, 1146 (15.0%) had two, 118 (1.5%) had three, 23 (0.3%) had four, and 8 (0.1%) had five or above. In Figure 7, we plot the genomic position of each detected breakpoint. We observe a recombination hotspot within spike.

**Figure 7.**
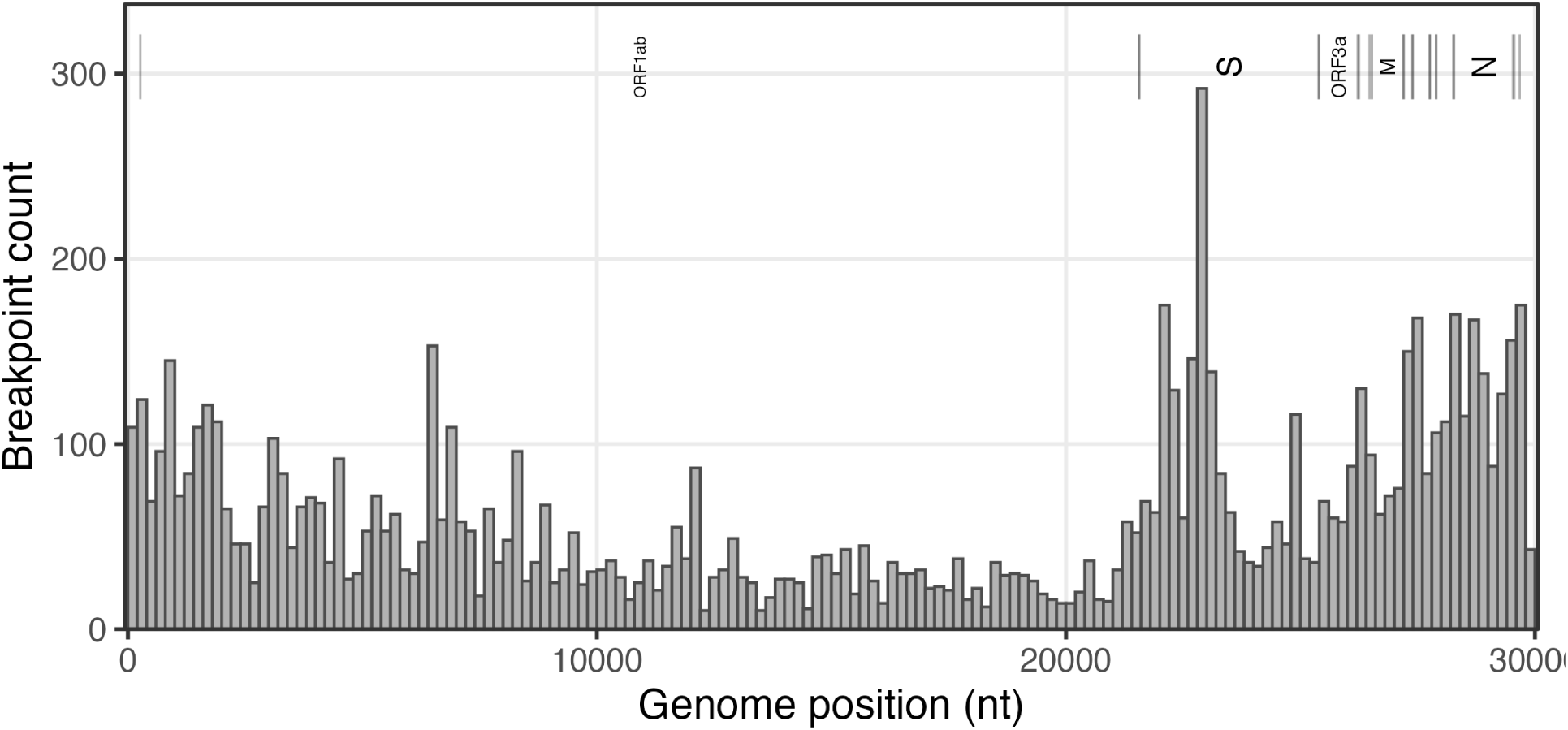
Histogram of detected recombination breakpoints.

We also observed enrichment of recombination breakpoints in intergenic regions. Gene boundaries were defined using the Wuhan-Hu-1 reference genome (GenBank: MN908947) (Benson et al. 2013; Wu et al. 2020). Although intergenic regions comprise only around 0.5% of the genome, 1.1% of all detected breakpoints were localized in these regions (90/8767). When calculating the proportion of detected breakpoints in intergenic regions, we excluded 339 breakpoints mapping to the ends of the genome, specifically from the 5’ end to ORF1ab and from ORF10 to the 3’ end. Using a two-sided binomial test, we found this enrichment to be highly statistically significant (*p* = 1.63 × 10^−9^).

## 4 Discussion

Genomic surveillance of recombinant SARS-CoV-2 sequences is important, given that mutations from each parental lineage can provide a growth advantage to the recombinant sequence. In this study, we developed an HMM to detect the local Pango lineage ancestry of query SARS-CoV-2 sequences based on lineage-specific nucleotide frequencies calculated using a reference set of recent sequences. Our method does not depend on an existing phylogeny, nor on any user-defined parameters such as the mutation rate or recombination rate. Instead, we use maximum likelihood to estimate the lineage-transition probability between consecutive sites, and the probability of observing alleles absent from the parental lineage at each site, which accounts for mutations on the recombinant sequence.

We validated our method using synthetic sequences generated from real SARS-CoV-2 genomes. In our simulation, our method achieved a sensitivity of 0.801 (95% CI: [0.775, 0.825]) for classifying recombinant sequences and a specificity of 0.989 (95% CI: [0.980, 0.994]) for classifying non-recombinant sequences. In 69.9% (95% CI: [67.0%, 72.7%]) of synthetic recombinant sequences, we detected two parental lineages that matched the true parental lineage pair. In 98.4% (95% CI: [97.4%, 99.1%]) of synthetic control sequences, we detected a single parental lineage that matched the true parental lineage. We found the sensitivity of our method to be positively associated with the number of mutations separating the parental sequences of the recombinant (see Figure 2). Finally, we estimated the mean distance between true and inferred breakpoints to be 1238 nucleotides (95% CI: [1108, 1386]) and 1007 nucleotides (95% CI: [901, 1125]) for synthetic recombinants with one and two breakpoints respectively.

Applying our model to real SARS-CoV-2 sequences collected in England between September 2020 and March 2024, we found 7619 recombinant sequences across 440,307 sequences, corresponding to 1.73% (95% CI: [1.69%, 1.77%]) of sequences. These 440,307 sequences were sampled across 185 test windows, each window corresponding to a 7-day period with no gaps between successive windows.

We hypothesized that across our test windows, the fraction of tested sequences detected as recombinant using our method would be positively associated with community SARS-CoV-2 prevalence, because higher prevalence raises co-infection opportunities, which should result in a higher rate of recombinant sequences in the population. We observed a positive association between the detected recombinant proportion in each test window and SARS-CoV-2 prevalence from the ONS survey, averaged within each test window (*p* = 0.0377).

However, we noted in Section 2.9 that our method can only detect recombinants whose parental sequences belong to distinct Pango lineages, and SARS-CoV-2 prevalence estimates do not consider the relative frequencies of circulating lineages. To address this, we also considered a lineage-adjusted measure of prevalence that incorporates the joint circulation of distinct Pango lineages, defined in equation S3 of the Supplementary Materials. We saw a strong positive association between the detected recombinant proportion in each test window and this lineage-adjusted measure of prevalence (*p* = 7.54 × 10^−5^). This finding indicates that higher detected recombinant proportions tend to coincide with windows in which the opportunity for co-infection between distinct Pango lineages is greater.

We next modeled the number of recombinants in our sample as a function of community SARS-CoV-2 prevalence to derive, for each parental lineage pair, the expected number of true positive recombinants across the 440,307 sequences analyzed (see equation S4 of the Supplementary Materials). For brevity, we refer to this as the expected true positive count for each parental lineage pair. This derivation relies on two key assumptions. First, we assume independent infections in our co-infection model, which means that the probability of co-infection by two lineages equals the product of their marginal prevalences. Second, we assume that the false positive rate and the detection factor (the product of sensitivity and the probability that a sequence from a co-infected individual is a recombinant) are constant across parental lineage pairs and test windows. We estimated the false positive rate and detection factor to be 0.011 (95% CI: [0.009, 0.013]) and 0.557 (95% CI: [0.297, 0.817]) respectively. Our estimated false positive rate closely matched the estimated false positive rate in our simulation study.

We estimated the expected true positive count for each parental lineage pair. We found that the number of detected recombinants with each parental lineage pair exceeded the corresponding expected true positive count for most parental lineage pairs. This is not surprising, because our expected true positive count does not include potential contributions from non-recombinant sequences that were detected as recombinant using our method. However, we cannot reliably allocate expected false positive counts across specific parental lineage pairs.

During the period when ONS SARS-CoV-2 prevalence estimates were available (until the test window ending March 19, 2023), we analyzed 387,054 sequences, with 5006 sequences detected as recombinant with two parental lineages. The vast majority of analyzed sequences should be non-recombinant. If our estimated false positive rate is correct, we would expect approximately 380, 000 × 0.01 = 3800 of these sequences to be false positive cases. Thus, the implied true positive count is approximately 5006 − 3800 = 1206 based on the estimated false positive rate. The expected true positive count summed across all lineage pairs was 1390. This shows that our estimated false positive rate and expected true positive counts are broadly consistent with the observed number of detected recombinants.

Although many detections are likely false positives, across parental lineage pairs, we found a strong positive association between the expected true positive count and the number of detected recombinants (*p* = 2.04 × 10^−6^). This indicates that our method is detecting recombinants at a rate that is predictable based on SARS-CoV-2 prevalence and co-infection dynamics between lineages under the null model of independent infections (Chin et al. 2024).

We then identified parental lineage pairs whose detected counts deviated from the overall trend between detected counts and expected true positive counts. We identified six under-represented parental lineage pairs (BA.1–BA.1.1, BA.1–BA.1.17.2, BA.1.1–BA.1.17.2, BQ.1.1–CH.1.1, BA.1.1–BA.2, CH.1.1–XBB.1.5). These lineage pairs, except for BA.1.1–BA.2, had lower detected counts than expected true positive counts.

Under-representation of BA.1–BA.1.1, BA.1–BA.1.17.2, and BA.1.1–BA.1.17.2 recombinants is likely explained by low sensitivity to detect these recombinants. Only a few mutations separate each of these lineage pairs, making recombinant detection difficult (see Figure 2).

On the contrary, pairwise Hamming distances are high for BQ.1.1–CH.1.1, BA.1.1–BA.2, and CH.1.1–XBB.1.5. However, if most of the genome is inherited from a single parent, the recombinant sequence can still be very similar to one of its parental lineages, so detection sensitivity may still be low. In Figure S2, we plotted the local Pango lineage ancestry of detected recombinants with parental lineage pairs BQ.1.1–CH.1.1, BA.1.1–BA.2, and CH.1.1–XBB.1.5. The lineage pairs BA.5.2–BA.5.2.1, BE.1.1–BQ.1.1, and AY.4–AY.4.2 have high detected and expected true positive counts and are included for comparison (see Figure 5). We found that detected breakpoints for lineage pairs BQ.1.1–CH.1.1, BA.1.1–BA.2, and CH.1.1–XBB.1.5 are more often at the ends of the genome relative to the other three lineage pairs with high detected counts, which would result in recombinant sequences with these parental lineage pairs that are indeed similar to one of their parental lineages. This could indicate that recombinants between these lineage pairs tend to have breakpoints near the ends of the genome, making their genomes similar to one of their parental lineages and resulting in low sensitivity to detect these recombinants.

Under-representation can also occur if parental lineage pairs were segregated to different geographical locations or subpopulations in England, which would make co-infection by these lineage pairs unlikely. Co-infection rates may be lower than expected even under homogeneous mixing, due to within-host interference. Moreover, lineage pairs may differ in their propensity to produce viable recombinants. Finally, the apparent under-representation of these parental lineage pairs could result from estimation error in expected true positive counts or sampling variability in detected counts. Estimating the variability of expected true positive counts is challenging. We do not have standard errors for ONS SARS-CoV-2 prevalence estimates. Additionally, this would require accounting for correlations in lineage prevalences across test windows.

We found that these six under-represented lineage pairs co-circulated in England during two intervals, late 2021 to early 2022 (BA.1–BA.1.1, BA.1–BA.1.17.2, BA.1.1–BA.1.17.2, BA.1.1–BA.2) and late 2022 to early 2023 (BQ.1.1–CH.1.1, CH.1.1–XBB.1.5). These two intervals coincide with periods when the estimated recombination proportion was low relative to ONS SARS-CoV-2 prevalence estimates. This indicates that these under-represented lineage pairs contributed to the lower frequency of recombinants detected during these periods. In particular, low sensitivity to detect recombinant sequences with parental lineage pairs BA.1–BA.1.1, BA.1–BA.1.17.2, and BA.1.1– BA.1.17.2 is likely contributing to the relatively low frequency of detected recombinants in late 2021 to early 2022.

Aggregating all detected recombination breakpoints, we observed a recombination hotspot within spike, which is consistent with previous work on recombination in SARS-CoV-2 and more broadly in sarbecoviruses (Lytras et al. 2022; Turakhia et al. 2022). Additionally, we found that breakpoints were enriched in intergenic regions, consistent with their high colocalization with TRS-B sites (Yang et al. 2021).

Using RIPPLES, Turakhia et al. (2022) found 2.7% of sampled genomes inferred to have detectable recombinant ancestry. This is higher than the proportion of detected recombinants using our method (1.73%; 95% CI: [1.69%, 1.77%]). This discrepancy is likely attributable to many factors. Turakhia et al. only analyze sequences up to May 2021, before the emergence of XBB. Furthermore, our method cannot detect recombination between sequences in the same lineage, which explains the lower proportion of detected recombinants using our method. Finally, even a modest difference in false positive rates would affect the estimated proportion.

Future work could estimate SARS-CoV-2 prevalence from the frequency of detected recombinants across test windows. In this study, we developed a statistical framework linking disease prevalence and lineage frequencies to the expected number of detected recombinants (see Section S2 of the Supplementary Materials). We further showed that these expected counts were correlated with observed recombinant counts. Estimating prevalence is feasible if the method’s sensitivity and false positive rate were known for the set of query sequences. In our study, we estimated the detection factor and false positive rate using ONS prevalence estimates (see Section S2 of the Supplementary Materials), so these rates are not generalizable outside England or beyond March 2023, when ONS prevalence estimates are no longer available. To estimate the prevalence of SARS-CoV-2 outside of England or beyond March 2023, we would need reliable sensitivity and specificity estimates for those populations and time periods.

Although we focused on SARS-CoV-2 in this study, our HMM is broadly applicable to other RNA and DNA viruses for detecting recombinants. Moreover, by limiting lineage transitions to predefined genome positions, our HMM can be easily adapted to detect reassortment events in segmented viruses such as influenza. Our detection method should also perform well on rapidly evolving viruses because we explicitly model novel alleles on recombinant sequences via a pseudo-frequency.

## 5 Implementation

The hidden Markov model and detection of recombinant sequences were implemented in Python 3.12.2. Results files were processed and plotted in R version 4.4.1.

## 6 Data and Resource Availability

All data and code for the analysis is available at github.com/nobuakimasaki/HMM-recombination. Portions of the preprocessing and analysis code were drafted with assistance from ChatGPT (GPT-4 and GPT-5). All AI-assisted code was reviewed and validated by the authors.

## Acknowledgments

This work is supported by NIH NIGMS R35 GM119774 to T.B. T.B. is a Howard Hughes Medical Institute Investigator.

We thank Professor Brian Browning for helpful suggestions on accelerating the forward algorithm and Professor Nicola Mueller for discussions of co-infection dynamics.

We gratefully acknowledge the investigators and laboratories that generated, submitted, and shared sequence data and metadata via GenBank (NCBI), which form the basis of this research.

