## Supplementary Materials for "Hidden Markov models detect recombination and ancestry of SARS-CoV-2"

### Supplementary information for *Hidden Markov models detect recombination and ancestry of SARS-CoV-2*

#### S1 Efficient forward algorithm

We implemented an efficient version of the forward algorithm, reducing the time complexity of the induction step from $\mathcal{O}(M^2)$ to $\mathcal{O}(M)$, where $M$ is the number of unique Pango lineages. Using the notation from the main paper, we define,

$$\alpha_t(i) = P(O_{1:t} = k_{1:t}, Z_t = i|\lambda, \epsilon),$$

which are our forward probabilities. This represents the probability of the observed nucleotide sequence up to position $t$ and the ancestral Pango lineage being lineage $i$ at position $t$.

In the induction step, we calculate the next time step for the forward probabilities. We have,

$$\alpha_{t+1}(j) = \left( \sum_{i=1}^{M} \alpha_t(i) a_{ij} \right) b_{j,t+1}(k_{t+1}).$$

Computing $\alpha_{t+1}(j)$ for one Pango lineage $j$ requires summing over $M$ lineages, which costs $\mathcal{O}(M)$. Thus, computing this for all Pango lineages costs $\mathcal{O}(M^2)$.

In our transition matrix, we have equal diagonal entries and equal off-diagonal entries. Recall,

$$a_{ij} = \begin{cases} 1-\lambda, & \text{if } i = j, \\ \frac{\lambda}{M-1}, & \text{if } i \neq j. \end{cases}$$

Furthermore, we use the scaled version of the forward probabilities, meaning that $\sum_{i=1}^{M} \alpha_t(i) = 1$. Thus, we can rewrite the induction step as,

$$\begin{aligned} \alpha_{t+1}(j) &= \left( (1-\alpha_t(j)) \frac{\lambda}{M-1} + \alpha_t(j)(1-\lambda) \right) b_{j,t+1}(k_{t+1}) \\ &= \left( \left( 1 - \lambda - \frac{\lambda}{M-1} \right) \alpha_t(j) + \frac{\lambda}{M-1} \right) b_{j,t+1}(k_{t+1}) \\ &= \left( \left( 1 - \frac{M}{M-1}\lambda \right) \alpha_t(j) + \frac{\lambda}{M-1} \right) b_{j,t+1}(k_{t+1}), \end{aligned}$$

which is constant time. Thus, computing this for all Pango lineages now costs $\mathcal{O}(M)$.

### S2 Expected recombinant counts

#### S2.1 Modeling lineage co-infection

We denote the lineage frequency of lineage $i$ in test window $w$, or the proportion of infections attributable to lineage $i$ among all SARS-CoV-2 infections during $w$, as $p_i(w)$. Note that $p_i(w)$ is distinct from the lineage prevalence, which is the overall fraction of the population infected by lineage $i$. We further denote the prevalence of SARS-CoV-2 in window $w$ as $\text{prev}(w)$. Then, it follows that the lineage prevalence of lineage $i$ in window $w$ is $\text{prev}(w)p_i(w)$.

Under the null model of independent infections described in Chin et al. (2024), the probability of co-infection by lineages $i$ and $j$ ($i < j$) is the product of their lineage prevalences. Thus,

$$P(\text{co-infected by } i \text{ and } j) = [\text{prev}(w)p_i(w)]\,[\text{prev}(w)p_j(w)] = \text{prev}(w)^2 p_i(w)p_j(w).$$

Conditioning on being infected (all of our sequences are from infected individuals) removes one factor of prevalence. We have,

$$P(\text{co-infected by } i \text{ and } j \mid \text{infected}) = \text{prev}(w)\, p_i(w)\, p_j(w).$$

Next, we denote the number of sequences for which we infer local Pango lineage ancestry in window $w$ as $n(w)$. In our study, $n(w)$ is known. If $X_{i,j}(w)$ denotes the number of sequences among $n(w)$ that are from individuals co-infected by lineages $i$ and $j$, it is reasonable that,

$$X_{i,j}(w) \sim \text{Binomial}\big(n(w), \text{prev}(w)\, p_i(w)\, p_j(w)\big).$$

##### S2.2 Modeling recombinant detection

However, not all sequences from individuals co-infected by lineages $i$ and $j$ will be $i$–$j$ recombinants. Furthermore, not all $i$–$j$ recombinants will be detected as recombinants using our method. To account for these two factors, we introduce two new parameters $\gamma_{i,j}(w)$ and $s_{i,j}(w)$:

$$\begin{aligned}
\gamma_{i,j}(w) &= P\left(\text{is } i\text{–}j \text{ recombinant} \mid \text{sequence from individual co-infected by } i \text{ and } j\right), \\
s_{i,j}(w) &= P\left(\text{detected as recombinant} \mid \text{is } i\text{–}j \text{ recombinant}\right).
\end{aligned}$$

Since $\gamma_{i,j}(w)$ and $s_{i,j}(w)$ are not separately identifiable, define the combined detection factor $\theta_{i,j}(w) = \gamma_{i,j}(w)s_{i,j}(w)$, and assume it is constant across lineage pairs and windows ($\theta_{i,j}(w) = \theta$).

Now let $R_{i,j}(w)$ denote the number of detected $i$–$j$ recombinants in window $w$. We have,

$$R_{i,j}(w) \mid X_{i,j}(w) \sim \text{Binomial}\big(X_{i,j}(w), \theta\big).$$

It follows that,

$$R_{i,j}(w) \sim \text{Binomial}\big(n(w), \text{prev}(w)\, p_i(w)\, p_j(w)\, \theta\big).$$

Taking the expected value,

$$\text{E}\left[R_{i,j}(w)\right] = n(w)\, \text{prev}(w)\, p_i(w)\, p_j(w)\, \theta.$$

However, we still have not considered the possibility that our method will classify non-recombinant sequences as recombinant. The above $R_{i,j}(w)$ is only comprised of true positive cases, but our method can also misclassify non-recombinant sequences as $i$–$j$ recombinants.

To account for this, we define,

$$\phi = P\left(\text{detected as recombinant} \mid \text{is non-recombinant}\right),$$

which is the false positive rate.

To simplify our derivations, we assume that $Y(w) = n(w) - \sum_{i<j} X_{i,j}(w)$ represents the number of sequences treated as non-recombinant in window $w$. Although this assumption does not hold when $\gamma_{i,j}(w) < 1$ for some $i < j$ (when not all sequences from individuals co-infected by $i$ and $j$ are $i$–$j$ recombinants), it is still reasonable because $n(w)$ is typically much larger than $\sum_{i<j} X_{i,j}$. This means that $Y(w) = n(w) - \sum_{i<j} X_{i,j}(w) \approx n(w) - \sum_{i<j} \gamma_{i,j}(w) X_{i,j}(w)$, with $n(w) - \sum_{i<j} \gamma_{i,j}(w) X_{i,j}(w)$ representing the correct number of non-recombinant sequences tested in $w$.

Then, denoting the set of sequences incorrectly classified as recombinants as $R^{FP}(w)$, we have that,

$$R^{FP}(w) \mid Y(w) \sim \text{Binomial}\left(Y(w), \phi\right).$$

Taking the expectation and using the tower rule,

$$\mathrm{E}\left[R^{FP}(w)\right] = \mathrm{E}\left[\mathrm{E}[R^{FP}(w) \mid Y(w)]\right] = n(w)\phi \left[1 - \text{prev}(w) \sum_{i<j} p_i(w) p_j(w)\right].$$

Then, denoting the total number of recombinants detected in $w$ as $R^{\text{total}}(w) = \sum_{i<j} R_{i,j}(w) + R^{FP}(w)$,

$$\begin{aligned}
\mathrm{E}\left[R^{\text{total}}(w)\right] &= n(w)\theta \text{prev}(w) \sum_{i<j} p_i(w) p_j(w) + n(w)\phi \left[1 - \text{prev}(w) \sum_{i<j} p_i(w) p_j(w)\right] \\
&= n(w) \left[\phi + (\theta - \phi)\, \text{prev}(w) \sum_{i<j} p_i(w) p_j(w)\right].
\end{aligned}$$

Thus,

$$\mathrm{E}\left[\frac{R^{\text{total}}(w)}{n(w)}\right] = \phi + (\theta - \phi)\, \text{prev}(w) \sum_{i<j} p_i(w) p_j(w). \tag{S1}$$

##### S2.3 Estimating detection parameters

We next want to estimate $\theta$ and $\phi$. Recall that $n(w)$ is known. We obtain our estimate of SARS-CoV-2 prevalence within window $w$, $\widehat{\text{prev}}(w)$, by averaging daily ONS SARS-CoV-2 prevalence estimates within window $w$. We obtain estimates for our lineage proportions,

$\hat{p}_i(w)$ and $\hat{p}_j(w)$, by taking the number of sequences that belong to lineage $i$ and $j$ respectively in window $w$, and dividing this by the total number of sequences in window $w$.

Let $x_w = \widehat{\text{prev}}(w) \sum_{i<j} \hat{p}_i(w)\hat{p}_j(w)$ and then equate $\text{E}\left[\frac{R^{\text{total}}(w)}{n(w)}\right]$ with $\frac{R^{\text{total}}(w)}{n(w)}$, which is the detected recombinant proportion in $w$ using our method described in the main paper. Now we have,

$$\frac{R^{\text{total}}(w)}{n(w)} = \phi + \big(\theta - \phi\big)x_w. \tag{S2}$$

We can estimate $\phi$ and $\theta$ by regressing $\frac{R^{\text{total}}(w)}{n(w)}$ on $x_w$. Denoting the fitted least squares intercept and slope as $\hat{\alpha}$ and $\hat{\beta}$ respectively, we calculate $\hat{\phi} = \hat{\alpha}$ and $\hat{\theta} = \hat{\alpha} + \hat{\beta}$.

#### S2.4 Expected true positive rate

Notice that in equation S1, the expected proportion of only true positive cases is,

$$\text{E}\left[\frac{\sum_{i<j} R_{i,j}(w)}{n(w)}\right] = \theta \text{prev}(w) \sum_{i<j} p_i(w)p_j(w).$$

Then, denoting our estimate of the expected true positive proportion in each window $w$ as $\hat{\rho}(w)$,

$$\hat{\rho}(w) = \hat{\theta}\widehat{\text{prev}}(w) \sum_{i<j} \hat{p}_i(w)\hat{p}_j(w). \tag{S3}$$

In our results section, we compare this with the detected recombinant proportion in window $w$ using our method described in the main paper. This detected proportion reflects the recombinants identified by our method and therefore includes both true positives and false positives. Thus, we expect the detected recombinant proportion to exceed $\hat{\rho}(w)$ in each window $w$. We are more interested in the correlation between these two quantities across windows.

Although $\theta$ is estimated using the detected recombinant proportions across windows, this does not constitute meaningful double use of the data. The regression is only used to estimate a single global scaling parameter $\theta$, assumed constant across all windows, and does not affect the strength of the linear association between $\hat{\rho}(w)$ and the detected recombinant proportion across windows.

#### S2.5 Expected counts by lineage pair

Finally, we obtain an estimate for the expected recombinant count by parental lineage pair. Recall that $R_{i,j}(w)$ represents the true positive $i$–$j$ recombinant count in $w$. Then, letting $\mathcal{W}$ be our set of test windows, we can denote the total true positive $i$–$j$ recombinant count across all windows as $R_{i,j} = \sum_{w\in\mathcal{W}} R_{i,j}(w)$. We have that $\mathrm{E}[R_{i,j}] = \theta \sum_{w\in\mathcal{W}} n(w)\mathrm{prev}(w)p_i(w)p_j(w)$. Then,

$$\hat{\mathrm{E}}[R_{i,j}] = \hat{\theta} \sum_{w\in\mathcal{W}} n(w)\widehat{\mathrm{prev}}(w)\hat{p}_i(w)\hat{p}_j(w). \tag{S4}$$

In our results section, we compare this estimate with the number of detected recombinants that have $i$ and $j$ as parental lineages. Because we do not allocate false positives across lineage pairs, $\hat{\mathrm{E}}[R_{i,j}]$ represents a true positive expectation, whereas the observed counts may include false positives. Thus, it is reasonable to expect that the observed count of $i$–$j$ recombinants will exceed $\hat{\mathrm{E}}[R_{i,j}]$. We are instead interested in the correlation between these two quantities across parental lineage pairs $i < j$.

#### S3 Supplementary Figures

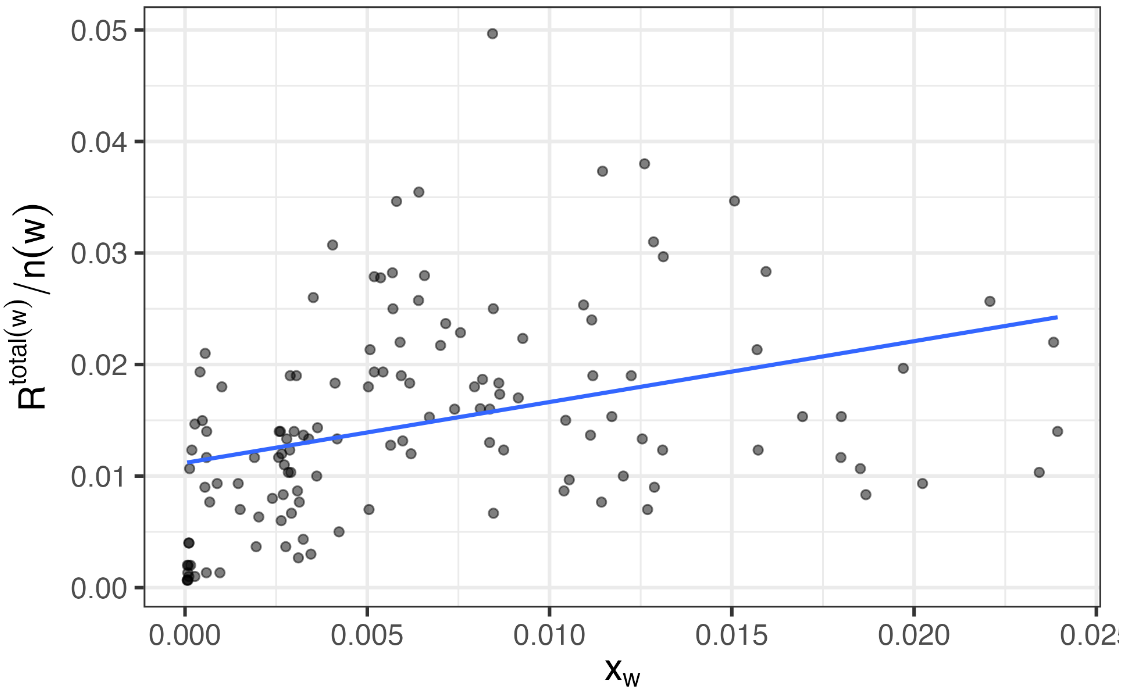


$\hat{\theta}$ = 0.557 (95% CI: [0.297,0.817]); $\hat{\phi}$ = 0.011 (95% CI: [0.009,0.013]); N = 132

**Figure S1.** Relationship between lineage co-circulation and detected recombinant proportion. Each point corresponds to a time window $w$. The horizontal axis shows $x_w = \widehat{\mathrm{prev}}(w)\sum_{i<j}\hat{p}_i(w)\hat{p}_j(w)$, where $\widehat{\mathrm{prev}}(w)$ is the window-averaged ONS SARS-CoV-2 prevalence and $\hat{p}_i(w)$ are estimated lineage proportions. The vertical axis shows the detected recombinant proportion $R^{\mathrm{total}}(w)/n(w)$. The fitted linear regression corresponds to $R^{\mathrm{total}}(w)/n(w) = \hat{\phi}+(\hat{\theta}-\hat{\phi})x_w$, with intercept $\hat{\phi}$ and slope $\hat{\theta}-\hat{\phi}$.

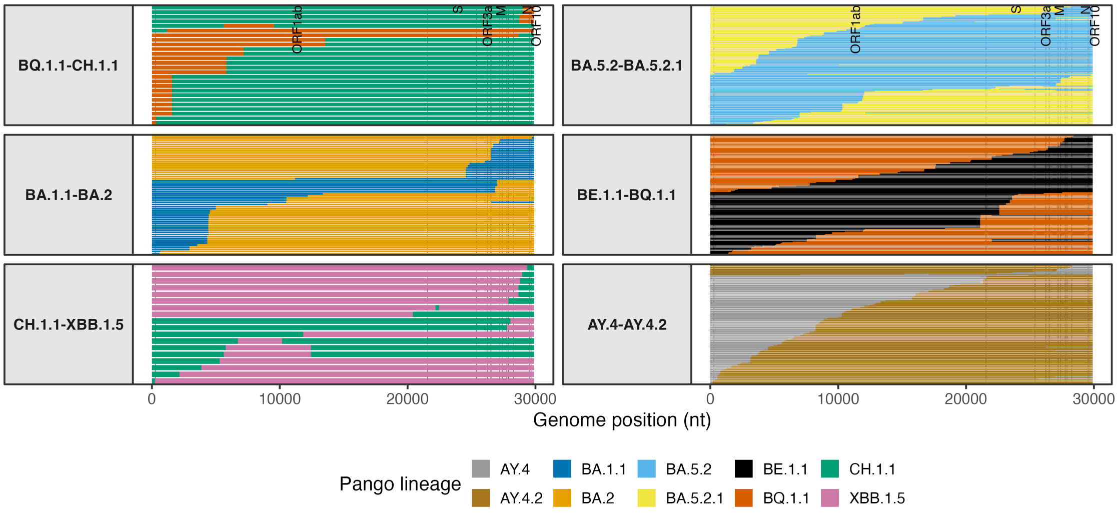


**Figure S2.** Local Pango lineage ancestry of recombinant sequences detected with lineage pairs BQ.1.1–CH.1.1, BA.1.1–BA.2, CH.1.1–XBB.1.5, BA.5.2–BA.5.2.1, BE.1.1–BQ.1.1, and AY.4–AY.4.2. Each row in each panel represents a detected recombinant sequence, with colors representing regions of the genome with distinct local Pango lineage ancestry. The lineage pairs BQ.1.1–CH.1.1, BA.1.1–BA.2, CH.1.1–XBB.1.5 are three of the under-represented lineage pairs, and the remaining lineage pairs are plotted for comparison. Detected sequences were filtered to the period in which ONS prevalence estimates were available.
